# Posterior axis formation requires *Dlx5/Dlx6* expression at the neural plate border

**DOI:** 10.1101/435321

**Authors:** Nicolas Narboux-Neme, Marc Ekker, Giovanni Levi, Églantine Heude

**Author notes:** Correspondence should be addressed to: Églantine Heude UMR7221 CNRS/MNHN 7, rue Cuvier 75231 Paris, Cedex 05, FRANCE.

## Abstract

Neural tube defects (NTDs), one of the most common birth defects in human, present a multifactorial etiology with a poorly defined genetic component. The *Dlx5* and *Dlx6* bigenic cluster encodes two evolutionary conserved homeodomain transcription factors, which are necessary for proper vertebrate development. It has been shown that *Dlx5/6* genes are essential for anterior neural tube closure, however their role in the formation of the posterior structures has never been described. Here, we show that *Dlx5/6* expression is required during vertebrate posterior axis formation. *Dlx5* presents a similar expression pattern in neural plate border cells during posterior neurulation of zebrafish and mouse. *Dlx5/6*-inactivation in the mouse results in a phenotype reminiscent of NTDs characterized by open thoracic and lumbar vertebral arches and failure of epaxial muscle formation at the dorsal midline. The *dlx5a/6a* zebrafish morphants present posterior NTDs associated with abnormal delamination of neural crest cells showing altered expression of cell adhesion molecules and defects of motoneuronal development. Our findings provide new molecular leads to decipher the mechanisms of vertebrate posterior neurulation and might help to gather a better understanding of human congenital NTDs etiology.

## INTRODUCTION

Neural tube defects (NTDs) correspond to a wide spectrum of common congenital disorders resulting from total or partial failure of neural tube closure during early embryogenesis. NTDs affect from 0.3 to 200 per 10 000 births worldwide ^1^ and vary in type and severity depending on the neural tube levels affected along the antero-posterior axis. Anterior and posterior neural tube defects lead respectively to brain (ie. exencephaly, anencephaly) or spinal cord malformations (ie. spina bifida); complete antero-posterior defect in neural tube closure is at the origin of a more severe form of NTD termed craniorachischisis (reviewed in ^2,3^). The origins of NTDs have been associated to genetic and/or environmental factors and more than 200 mutant mice have been reported to present different forms of neural tube malformations ^4,5^. However, given the complexity of the NTD spectrum, there has been limited progress in defining the molecular basis of these conditions.

In vertebrates, neural tube defects originate from a failure in morphogenetic events taking place during the neurulation process. In mammalian embryos, neurulation involves two distinct morphogenetic processes along the rostro-caudal axis, known as primary and secondary neurulations. Primary neurulation refers to neural tube formation originating from folding of an open neural plate that forms the central lumen in the anterior part of the embryo. In contrast, secondary neurulation is characterized by mesenchymal condensation and cavitation in the posterior axis caudal to the tail bud ^6,7^. In zebrafish, neurulation occurs homogeneously along the rostro-caudal axis by epithelial condensation forming the neural plate followed by cavitation as observed during secondary neurulation in mammals ^8,9^. Zebrafish neurulation has been linked either to primary or to secondary neurulation of higher vertebrates ^6,7^. However, the morphogenetic similarities observed between neurulation in teleosts and other vertebrates indicate that zebrafish neural tube formation rather correspond to primary neurulation and constitutes a viable model to study vertebrate neural tube development ^6,10^.

The general primary neurulation dynamic seems conserved among vertebrates and is characterized by convergent movement of the neural plate borders (NPB) toward the dorsal midline to generate the neural tube with a central lumen ^6,9^. NPB cells constitute a competence domain, established between neural and non-neural ectoderm, that delineates the presumptive domain at the origin of migratory neural crest cells (NCCs) and responsible for neural tube closure ^11,12^.

*Dlx* genes, the vertebrate homologues of *distal-less (dll)* in arthropods, code for an evolutionary conserved group of homeodomain transcription factors. The mouse and human *Dlx* gene system is constituted by three closely associated bigenic clusters located on the same chromosomes as *Hox* genes clusters. In teleost, the *dlx* clusters are arranged on chromosomes similarly to their tetrapod *Dlx* counterparts ^13^. The most probable scenario suggests that *Dlx* genes have arisen from an ancestral *dll* gene as a result of gene duplication events ^14^. Data indicate that *Dlx* genes from a same cluster, such as *Dlx5* and *Dlx6* paralogs, present redundant functions during vertebrate development ^15-18^.

It has been shown that *Dlx5* is one of the earliest NPB markers defining the limit of the neural plate during neurulation of mouse, chick, frog and zebrafish ^18-24^. Inactivation of *Dlx5* in mouse results in a frequent exencephalic phenotype suggesting defects of anterior neural tube closure ^16^, however the mice do not present obvious posterior axis malformations. As *Dlx5* and *Dlx6* have partially redundant functions, it has been necessary to simultaneously inactivate both genes to fully reveal their roles during development. Functional analyses of *Dlx5/Dlx6* inactivation in mice, avian and fish have demonstrated evolutionary conserved roles in appendage morphogenesis, in neurogenesis, in the development of the face and of the reproductive system ^16-18,25-31^. *Dlx5/6*^-/-^ mice also present midline-fusion abnormalities including hypospadias, failure of anterior neuropore closure and posterior axis malformations ^17,31,32^. However, the origin of the latter phenotype has never been described.

Here we show that simultaneous invalidation of *Dlx5* and *Dlx6* in zebrafish and mouse results in defects of posterior axis formation. Our data indicate a conserved role for *Dlx5/6* in posterior neurulation in vertebrates and suggest that genetic pathways involving these genes might be implicated in syndromic forms of human midline defects.

## RESULTS

### *Dlx5/6* invalidation induces posterior axis malformations in zebrafish and mouse

To analyse the effect of *Dlx5/6* invalidation on axis formation, we generated *dlx5a/6a* zebrafish morphants and *Dlx5/6*^-/-^ mouse embryos and compared the resulting axial phenotypes. The disruption of *Dlx5/6* function led to similar early posterior malformations characterized by curly-shaped tails in both species (Fig. 1 A-D, white arrowheads). In *Dlx5/6* mutant mice, 80% of embryos and foetuses presented a curly-shaped tail associated with varying degrees of exencephaly (Fig. 1, blue arrowhead), the latter phenotype known to result from defect of anterior neural tube closure ^17,19,25,33^. Moreover, E18.5 mutant mice displayed medio-dorsal split in the thoracic/lumbar region (Fig. 2A, B, red arrowheads). Skeletal preparations and immunostainings on sections revealed lack of vertebral arches fusion dorsally at both thoracic and lumbar levels and failure of epaxial muscle formation at the dorsal midline (Fig. 2 C-F, red arrowheads).

**Figure 1.**
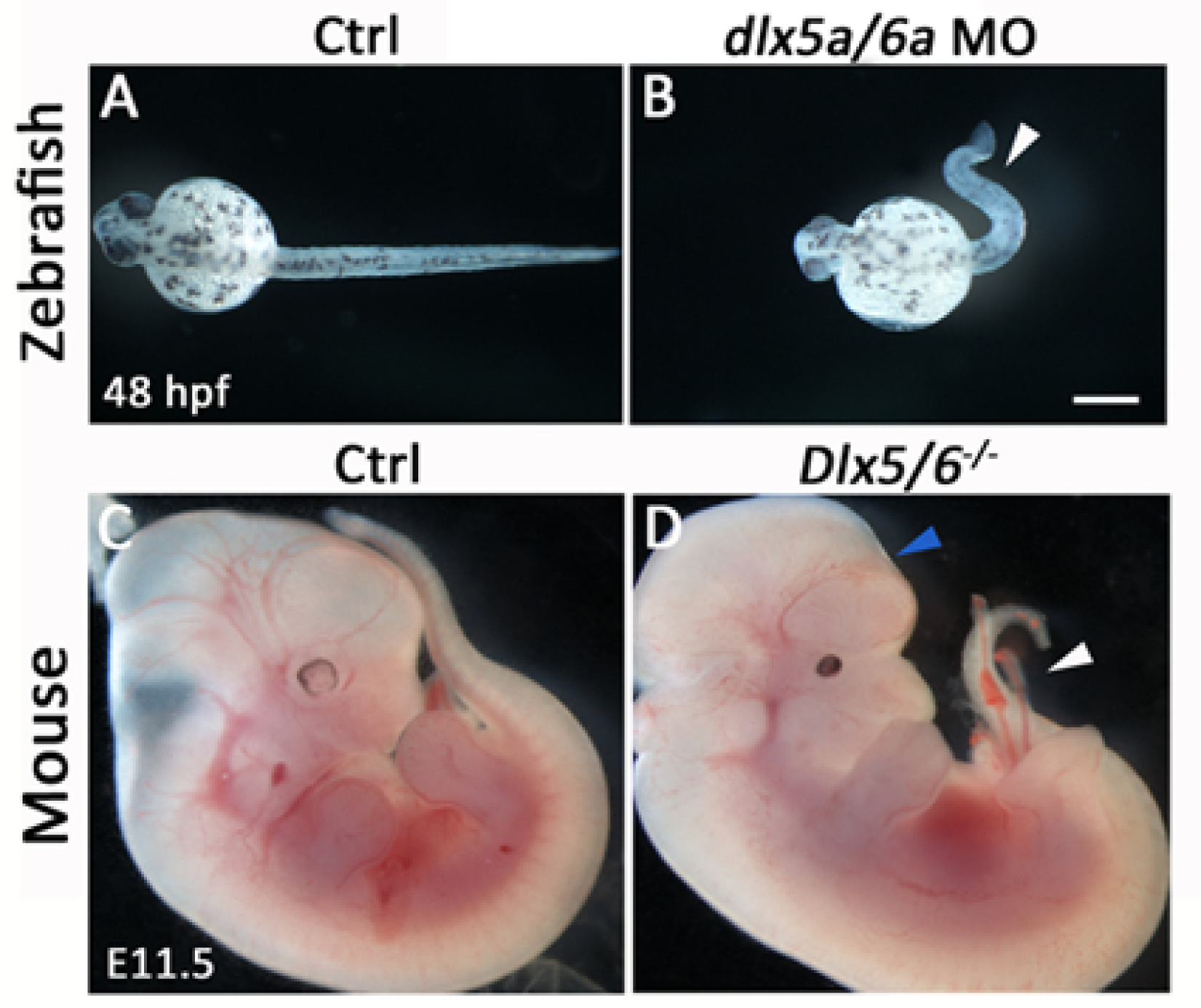
Early phenotype of *Dlx5/6*-invalidated zebrafish and mouse. (A-B) Phenotype of a control zebrafish and a *dlx5a/6a* morphant at 48 hpf. (C-D) Phenotype of a control and a *Dlx5/6*^*-/-*^ mutant mouse at E11.5. Invalidation of *Dlx5/6* in zebrafish and mouse leads to early defect of axis development characterized by curly-shaped tail phenotype in both models (B, D, white arrowheads). In *Dlx5/6*^*-/-*^ embryos, the caudal phenotype is associated with defect of brain formation (D, blue arrowhead). Scale bar in B for A-B 100μm, for C-D 1000μm.

**Figure 2.**
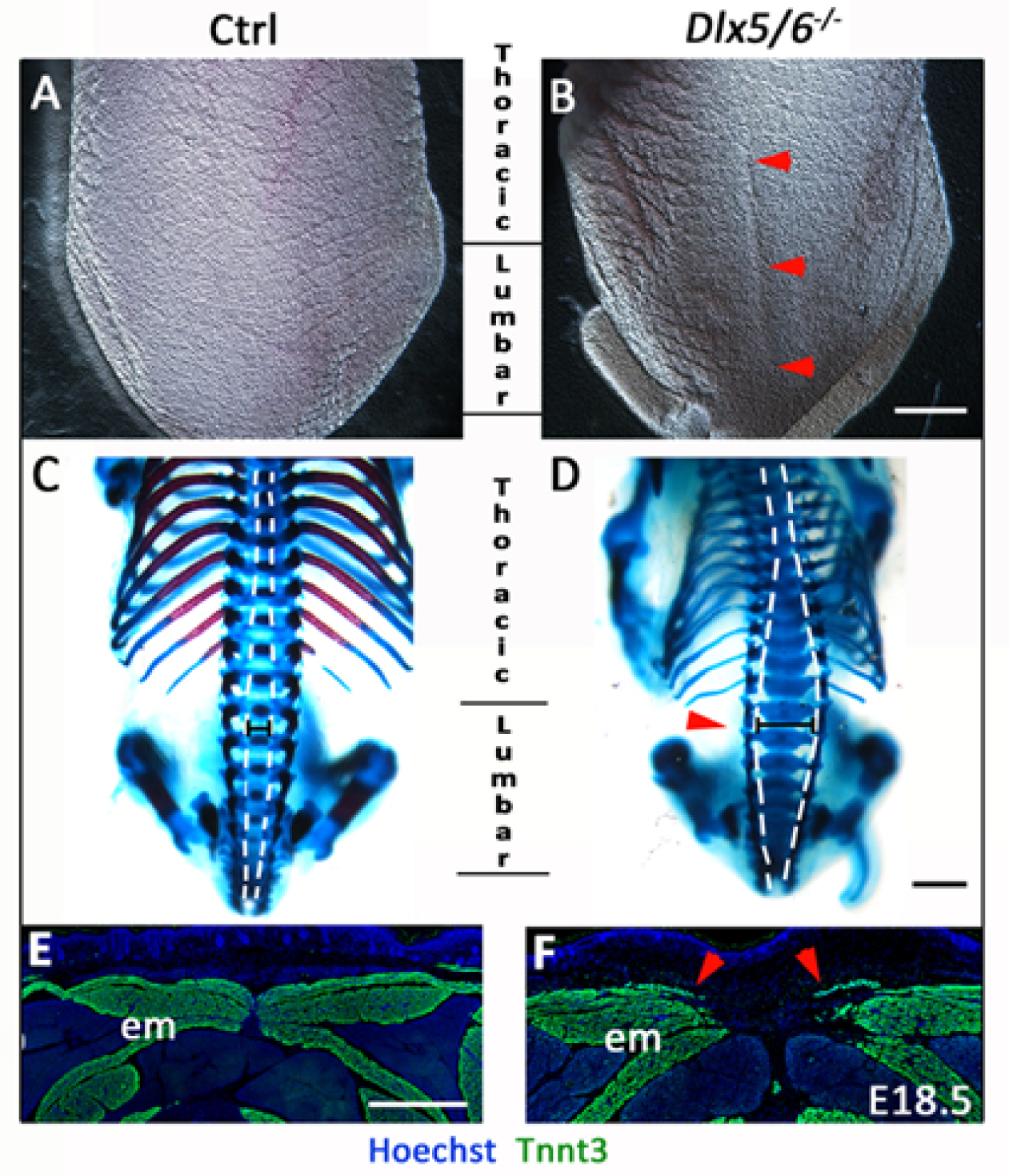
Dorsal midline defects in perinatal *Dlx5/6*^*-/-*^ mice. (A-D) Macroscopic dorsal view (A-B) and skeletal preparation (C-D) of the posterior axis of control and *Dlx5/6*^*-/-*^ mutant mice at E18.5. (E-F) Immunostaining on coronal cryosections for Tnnt3 in dorsal musculature of control and *Dlx5/6*^*-/-*^ E18.5 foetuses. The *Dlx5/6*^*-/-*^ mutants display apparent dorsal split associated with defects of thoracic/lumbar vertebrae and of epaxial muscle formation at the midline (B, D, F red arrowheads). Abbreviations: epm, epaxial muscles. Scale bars in B for A-B, in D for C-D 2000μm and in E for E-F 200μm.

### Expression of *Dlx5* during zebrafish and mouse posterior axis formation

To understand the origin of the axis phenotype observed in *Dlx5/6*-invalidated specimens, we then compared the spatio-temporal expression of *Dlx5* during zebrafish and mouse posterior neurulation. We previously showed in zebrafish that *dlx5a*-expressing ectodermal cells are laterally connected to the neural ectoderm to form the presumptive median fin fold ^18^. At 15.5 hpf, *dlx5a*-expressing NPB cells along the neural keel follow a medial convergence toward the dorsal midline during neural rod formation at 16 hpf (Fig. 3A-B, black arrowheads). At later stages, *dlx5a* expression is limited to median fin fold ectodermal cells at 24 hpf and 48 hpf, and gradually decreased until 72 hpf ^18^.

**Figure 3.**
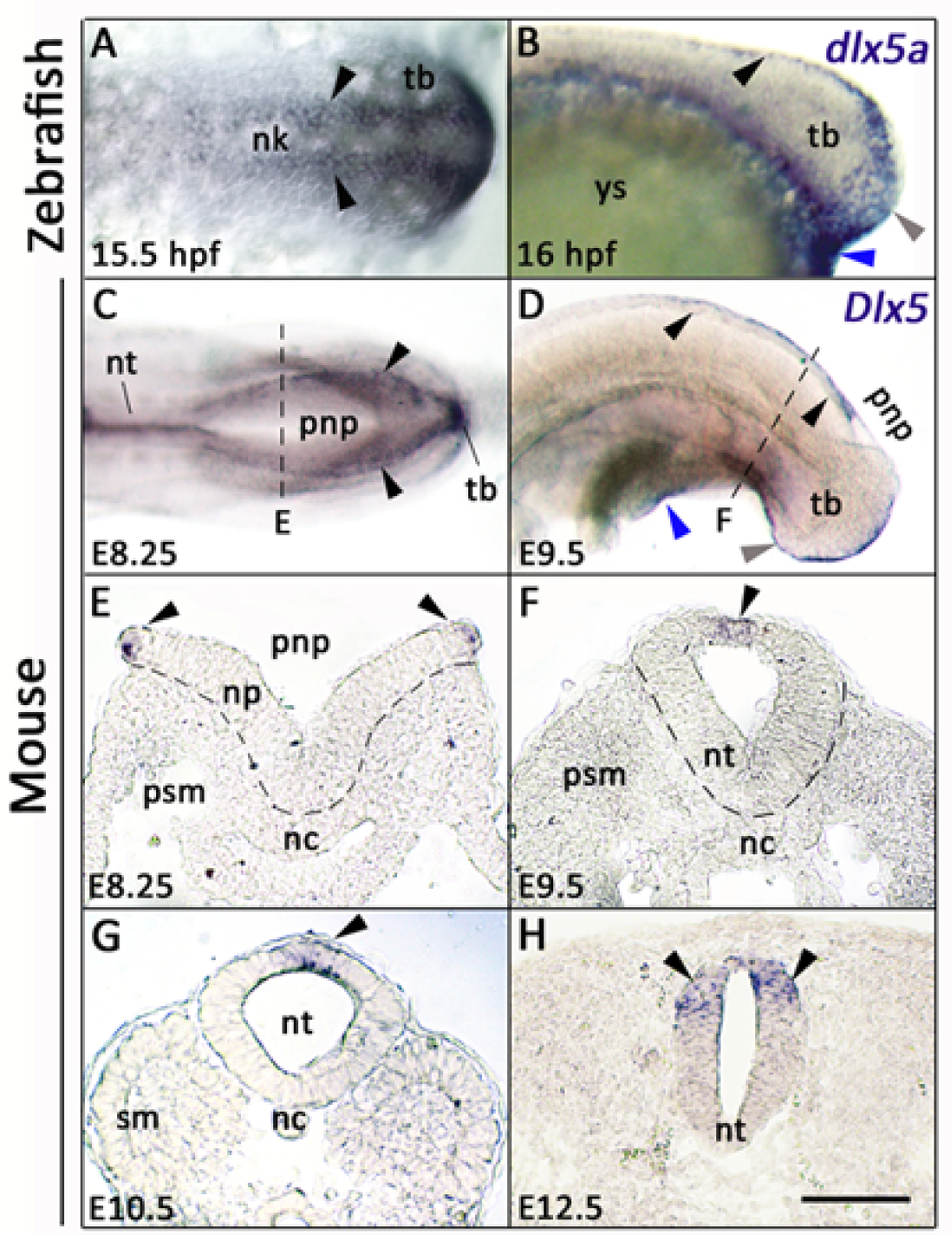
Expression analysis for *Dlx5* during zebrafish and mouse posterior neurulation. (A-D) Whole-mount *in situ* hybridization for *dlx5a* and *Dlx5*; (A, C) dorsal and (B, D) lateral views of the posterior axis of 15.5 hpf and 16 hpf zebrafish (A-B) and E8.25 and E9.5 mouse embryos (C-D). (E-F) *In situ* hybridization for *Dlx5* on coronal cryosections at the levels indicated by the dashed lines in (C, D and Supplementary Fig. S1 online). In zebrafish embryos, *dlx5a* transcripts are detected in NPB cells along neural keel at 15.5 hpf and at the dorsal midline at 16 hpf (A-B, black arrowheads). During mouse posterior neurulation, *Dlx5* is expressed in NPB cells surrounding the posterior neuropore and along the dorsal midline of the neural tube after neural tube closure (C-H, black arrowheads). In both species, *Dlx5* is also detected in the ventral ectodermal ridge of the tail bud and at the cloacal level (grey and blue arrowheads respectively in B, D). Abbreviations: nc, notochord; nk, neural keel; np, neural plate; nt, neural tube; pnp, posterior neuropore; psm, presomitic mesoderm; sm, somitic mesoderm; tb, tail bud; ys, yolk sac. Scale bar in H for A, H 50μm, for B, E-G 75μm, for C 150μm, for D 200μm.

Similarly, in mouse embryos, *Dlx5* transcripts were detected in NPB cells surrounding the posterior neuropore and at the dorsal midline after neural tube closure at E8.25 and E9.5, with gradual decrease of expression in a rostro-caudal manner (Fig. 3C-F, black arrowheads). At E10.5 and E12.5, *Dlx5* expression was maintained in the dorsal neural tube after posterior neuropore closure (Fig. 3G-H, black arrowheads). In both models, we also observed *Dlx5* expression in the ventral ectodermal ridge (VER) of the tail bud and at the cloacal level (Fig. 3B, D; Supplementary. Fig. S1 online, grey and blue arrowheads respectively). The expression and functional analyses in zebrafish and mouse suggest a conserved role of *Dlx5/6* genes in posterior axis development.

### Posterior neurulation defects in *dlx5a/6a* zebrafish morphants

To further investigate the role of *Dlx5/6* genes during posterior neurulation, we next performed molecular analyses in early *dlx5a/6a* zebrafish morphants during neural keel-rod transition at 16 hpf. Given the key role for cell adhesion molecules (CAM) in neural tube morphogenesis, neural tube closure and epithelial-to-mesenchymal transition ^10,34-37^, we analysed the expression of *ncad* (*cdh2*) and *ncam3*, members of the CAM family involved in cell-cell adhesion. We also analysed expression of *msx1b*, marker of NPB cells and premigratory neural crest cells (NCCs) ^38,39^, and expression of *foxd3*, marker of premigratory and early migratory NCCs ^40^.

In 16 hpf control embryos, *ncad* is constitutively expressed in the neural keel and the presomitic/somitic mesoderm. In contrast, *ncam*3 expression is limited to the dorsal part of the neural keel (Fig. 4A-A’, C-C’). In *dlx5a/6a* morphants, we observed a loss of *ncad* and *ncam*3 expression in aberrant protruding cells at the dorsal midline of the neural keel (Fig. 4B’, D’, black arrowheads). Moreover, whole-mount expression pattern of *msx1b* and *foxd3* in morphants revealed a defect of neural tube formation compared to controls, with bifid stripes of expression in the caudal-most part of the axis, characteristic of a delay in neural keel-rod transition (Fig. 4F-H, grey arrowheads). On sections, *msx1b* showed a decrease of expression at the midline where protruding cells were detected in morphants (Fig. 4E’, F’, black arrowhead). At 16 hpf, *foxd3* expression in the dorsal neural tube labelled premigratory NCCs (Fig. 4G’). The analysis of *dlx5a/6a* morphants revealed that the aberrant protruding cells at the dorsal midline of the neural keel were positive for *foxd3* expression (Fig. 4H’, black arrowhead). The results indicated that disrupted *dlx5a/6a* function affects *msx1b* and CAM (*ncad/ncam3*) expression at the roof plate of the neural keel in aberrant delaminating *foxd3*-positive NCCs.

**Figure 4.**
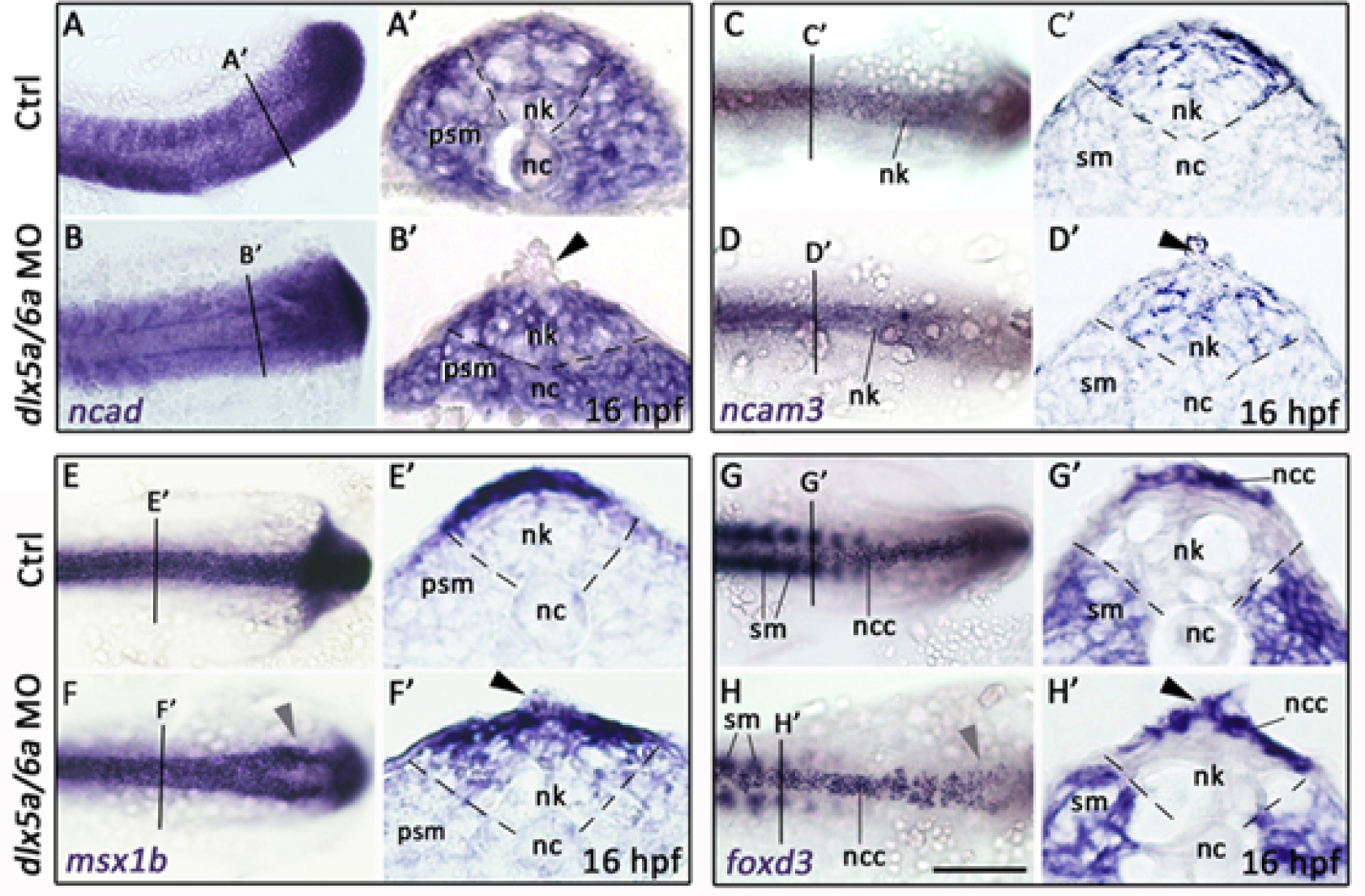
Altered early posterior neurulation in *dlx5a/6a* zebrafish morphants. (A-H) Dorsal views of whole-mount *in situ* hybridization for *ncad, ncam3, msx1b* and *foxd3* in controls and *dlx5a/6a* morphants at 16 hpf. (A’-H’) *In situ* hybridization for *ncad, ncam3, msx1b* and *foxd3* on coronal cryosections of 16 hpf controls and *dlx5a/6a* morphants at levels indicated by lines in (A, H). During neural keel-rod transition, the *dlx5a/6a* morphants show a decrease or loss of *ncad, ncam3* and *msx1b* in aberrant protruding *foxd3*-positive NCC at the dorsal midline of the neural keel (black arrowhead in B’-H’). Abbreviations: nc, notochord; ncc, neural crest cells; nk, neural keel; psm, presomitic mesoderm; sm; somitic mesoderm. Scale bar in H for A-H 100μm, for A’-H’ 25μm.

We also studied the effect of *dlx5a/6a* invalidation at later stages when the neural tube is formed. In 24 hpf controls, dorsal *foxd3* expression in migratory NCCs was observed in the caudal neural tube (Fig. 5A-A’). In *dlx5a/6a* morphants, *foxd3*-positive cells showed abnormal asymmetric profile (Fig. 5B’, black arrowhead) associated with a reduced neural tube and defect of lumen formation (Fig. 5A’, B’, dashed lines), the latter phenotype being well revealed by the constitutive *ncad* expression in the neural tissue (Fig. 5C’, D’). The *foxd3* and *ncad* transcripts were also detected in the somitic mesoderm of controls, that later develops into myotomes arranged into V-shaped chevrons (Fig. 5A, C, Supplementary. Fig. S2 online). In *dlx5a/6a* morphants, *foxd3* and *ncad* expression patterns revealed defect of somitic segmentation at 16 hpf and abnormal U-shaped chevrons at 24 hpf (Fig. 4A-B, G-H; Fig. 5A-D). The expression profile of *msx1b* along the neural tube was also altered in *dlx5a/6a* MO embryos at 24 hpf (Fig. 5E-F, black arrowhead), while expression in the median fin fold compartment seemed unaffected.

**Figure 5.**
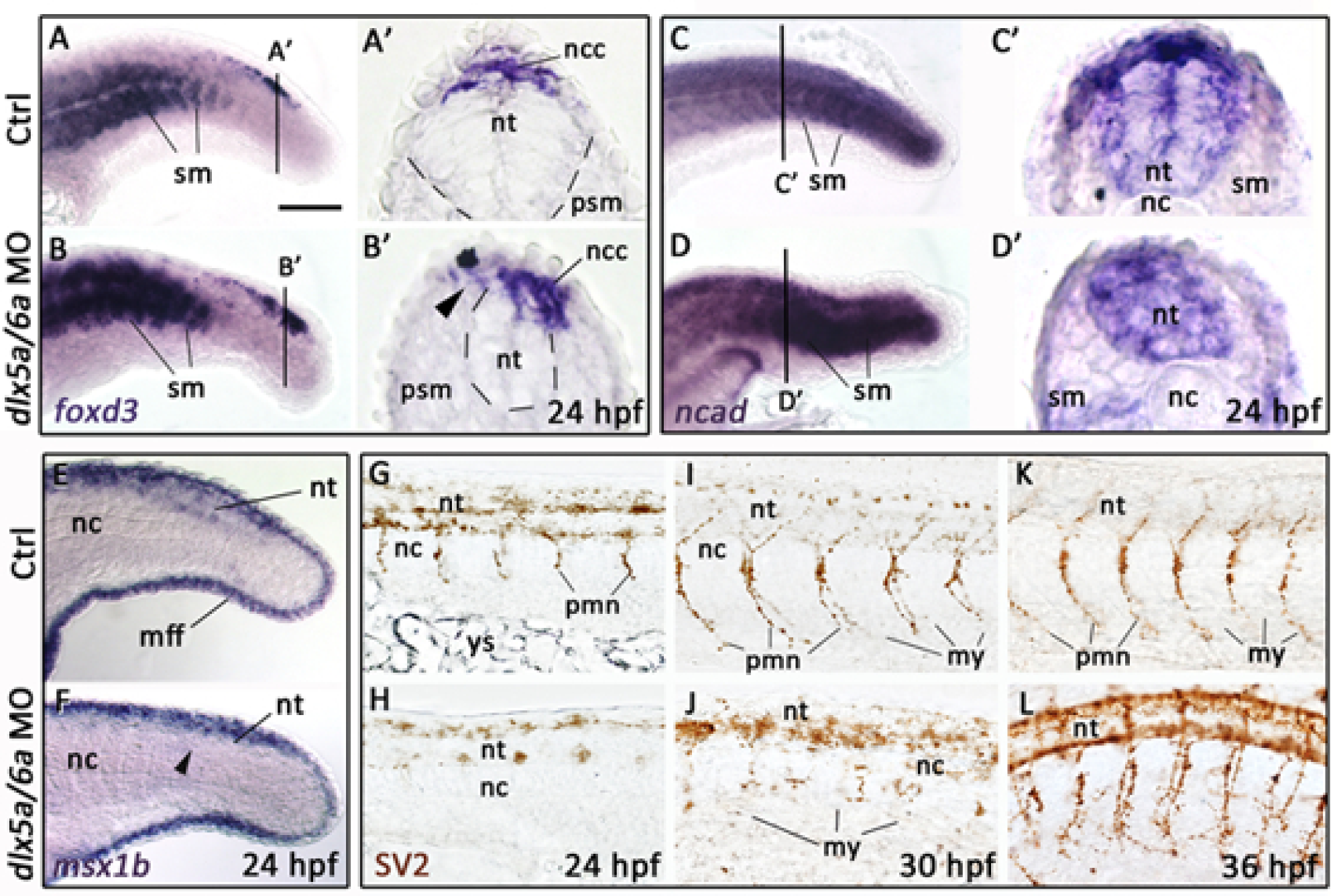
Defects of neural tube and motoneuron formation in *dlx5a/6a* zebrafish morphants. (A-F) Lateral views of whole-mount *in situ* hybridization for *foxd3, ncad* and *msx1b* in controls and *dlx5a/6a* zebrafish morphants at 24 hpf. (A’-D’) *In situ* hybridization for *foxd3* and *ncad* on coronal cryosections at levels indicated by lines in (A-D). The *dlx5a/6a* morphants show a reduced neural tube associated with a defect of NCC migration, abnormal somitic segmentation and decrease of *msx1b* in the neural tube (A-F). (G-L) Lateral views of whole-mount immunostaining for SV2 in 24 hpf, 30 hpf and 36 hpf control and *dlx5a/6a* morphant embryos. In *dlx5a/6a* morphants at 24 hpf and 30 hpf, axonal projections of primary motoneurons fail to form (G-J). At 36 hpf, primary motoneurons show aberrant synaptic connection to their myotomal targets in *dlx5a/6a* morphants (K, L). Abbreviations: mff, median fin fold; my, myotomes; nc, notochord; ncc, neural crest cells; nt, neural tube; pmn, primary motoneurons; psm, presomitic mesoderm; sm; somitic mesoderm, ys, yolk sac. Scale bar in A for A-F 100μm, for A’-D’ 25 μm and for G-L 50μm.

We next analysed in *dlx5a/6a* morphants the primary motoneuron (PMN) population, originating from NCCs and known to be affected in NTDs ^41,42^. In 24 hpf controls, the PMNs express the synaptic vesicle SV2 and axonal projections start to elongate from the neural tube to reach their myotomal targets, forming the neuromuscular junctions at 30 hpf and 36 hpf (Fig. 5G, I, K). In contrast, *dlx5a/6a* MO showed defect of motoneuronal outgrowth at 24 hpf and PMNs failed to connect the myotomes at 30 hpf (Fig. 5H, J). At 36 hpf, the axonal projection was completed but PMNs showed defect of neuromuscular junction with aberrant synaptic arborescence (Fig. 5L). Moreover, defect of PMN formation was associated with accumulation of SV2 protein in the dorsal neural tube at 30 hpf and 36 hpf (Fig. 5J, L). The defect of neuromuscular innervation was also observed in *Dlx5/6* mutant mice as shown by immunostainings on sections for the neuronal marker Tuj1 in trunk epaxial muscles positive for Tnnt3 (Supplementary. Fig. S3 online, white arrowheads).

We also studied the expression of *shha* and *bmp4* that are organizers of tail development ^11,43-46^. In 24 hpf controls, *shha* is expressed in the notochord, the neural tube floor plate and in the tail stem cell pool, namely the chordoneural hinge (Supplementary. Fig. S2 online). At 48 hpf, *shha* expression is limited to the floor plate (Supplementary. Fig. S2 online). In morphants, *shha* expression was maintained but well reveals the undulating phenotype of axial structures (Supplementary. Fig. S2 online). While *bmp4* did not show obvious defect of expression at 16 hpf and 24 hpf ^18^, expression in the spinal cord was altered at 48 hpf (Supplementary. Fig. S2 online).

Altogether, our data indicated that disrupted *dlx5a/6a* function in zebrafish led to a loss of cell adhesion molecules expression in protruding NCCs at the dorsal midline, resulting in posterior neural tube defects and mispatterned NCC-derived primary motoneurons.

## DISCUSSION

It has been described previously that *Dlx5* is one of the earliest NPB marker during gastrulation ^20,21,24^. Particular attention has been paid on its role during anterior neural tube formation in defining the border between non-neural and neural plate territories ^11,21,33,47,48^. However, the role of *Dlx5/6* during posterior neurulation has never been reported.

Our analysis highlights the role of *Dlx5/6* genes in NPB cells during posterior neurulation in zebrafish and mouse (Figs. 1-5) ^18^. In both models, *Dlx6* expression is undetectable by *in situ* hybridization during early neurulation. In zebrafish, *dlx6a* expression is observed in the presumptive median fin fold that is related to *dlx5a*-expressing NPB cells during unpaired fin development ^18^. *Dlx6* expression appears delayed and weaker compared to expression of the *Dlx5* paralog, a difference that was observed by others and us in vertebrate embryos using a variety of probes for these genes. The invalidation of both genes is however required to observe a fully penetrant phenotype, underlying the redundant functions of *Dlx* paralogs in vertebrates ^16-19,49-51^.

Expression analyses for *Dlx* homologs in various models including chick, xenopus, lamprey and amphioxus suggest that *Dlx* expression in NPB cells has been established early during chordate evolution ^21,24,47,52,53^. However, we can still notice differences in *Dlx5* expression along the rostro-caudal axis of zebrafish and mouse. In mouse, *Dlx5* is expressed in NPB cells during both anterior and posterior neurulation (Fig. 3) ^20^. In amphioxus, *amphiDll* is expressed in NPB cells all along the rostro-caudal axis, suggesting a conserved role of *Distal-less-*related genes in anterior and posterior neurulation in chordates ^53^. In contrast, in zebrafish, the anterior limit of *dlx5a*-expressing NPB cells is located at the 8^th^ somite level and appears closely related to the establishment of the presumptive median fin fold (Fig. 3) ^18^. According to our expression analysis, we did not find evidence of anterior neurulation defects in *dlx5a/6a* morphants ^29^. In teleosts, neurulation is characterized by uniform epithelial condensation and cavitation, which give rise to both anterior and posterior neural tube ^6,8^. It has been suggested that zebrafish might present primitive mechanisms of neurulation, however basal chordates show primary and secondary neurulation as observed in higher vertebrates ^6,54^. Our results indicate that, while the cellular morphogenetic basis of neural tube formation is uniform along the rostro-caudal axis of zebrafish, the anterior and posterior neurulation processes do not involve same molecular mechanisms, suggesting evolutionary divergence of *Dlx5/6* function during anterior neural tube formation in teleosts. Our data bring new insights into the genetic and evolutionary origins of neural tube formation in chordates. Special attention should be paid in future studies to elucidate the genetic requirement differences between anterior and posterior neurulation in teleosts.

Our analysis reveals that disrupted *Dlx5/6* function in zebrafish and mouse leads to early curly-shaped tail phenotypes in both models (Fig. 1). In *Dlx5/6*^-/-^ mice, the tail phenotype is associated with midline axis defects and brain malformations characteristic of NTDs (Figs. 1-2) ^17,19,25^. Intriguingly, the axis defects observed in *Dlx5/6*^*-/-*^ mice was similar to the phenotype of *CT “curly tail”* mutants, a historical model of NTDs ^5,55,56^. The data thus suggested that the axis phenotype observed in *Dlx5/6*-invalidated zebrafish and mice resulted from defects of posterior neurulation, an aspect that we confirm through our functional analysis in zebrafish.

We show that *dlx5a/6a* morphants present neural tube defects associated with aberrant dorsal delamination and migration of NCCs (Figs. 4-5). The protruding NCCs observed at the dorsal midline of the neural keel show loss of cell adhesion molecule transcripts (Fig. 4). It has been shown that these latter are important actors during neurulation ^7^. In particular, *ncad* is required for NPB convergence, neural tube closure, maintenance of neural tube integrity and epithelial-to-mesenchymal transition ^10,34-37^. In zebrafish, *ncad* invalidation represses neural tube formation due to defect of convergence and intercalation of NCCs ^10,34,35^. The data thus indicate that *dlx5a/6a* genes act in NPB cells for adhesion integrity of NCCs during neural tube formation.

We also noticed that zebrafish morphants present failure of somite segmentation and myotomal morphology (Figs. 4-5) that might indirectly originates from NTDs or defect of signalling from the VER. The latter is known to act as a signalling centre during tail somitogenesis and elongation ^57,58^. VER cells undergo epithelial-to-mesenchymal transition during tail development as observed dorsally in NCCs ^59^. The continuous dorso-ventral ectodermal *Dlx5*-positive domain might reflect that the VER represents a ventral extension of dorsal ectodermal cells during tail morphogenesis. The inhibition of BMP signalling by *Noggin* suppresses the VER epithelial-to-mesenchymal transition process during tail morphogenesis ^57,59^. In *dlx5a/6a* morphants, ectodermal *Bmp4* expression is not affected during early stage of posterior axis development ^18^, and the expression of *Noggin* has been shown to be independent of *Dlx5* during craniofacial development ^60^. *Dlx5/6* expression in the VER does not appear required for tail elongation as both *Dlx5/6*^-/-^ mice and *dlx5a/6a* morphants show equivalent number of axial segments compare to controls. However, the defect of somitic mesoderm patterning in *dlx5a/6a* morphants (Fig. 5) suggests a potential role for *Dlx5/6* in NPB and/or VER cells during tail segmentation.

Taken together, our findings reveal the role of *Dlx5/6* genes in vertebrate posterior axis formation. It has been demonstrated previously that *Dlx5* acts to specify NPB cells during cranial neural tube formation in mouse and chick ^20,21,24^. However, *Dlx* activity in xenopus is not necessary for induction of NCCs ^47^. Our findings confirm that *Dlx* invalidation does not impact on NCC induction as *foxd3* expression is maintained in the protruding NCCs observed in *dlx5a/6a* morphants (Fig. 4). In addition, *msxb1* expression is decreased in the protruding NCCs of *dlx5a/6a* morphants, associated with defects in primary motoneurons projections (Figs. 4-5). It has been shown that *msx* and *ncad* genes are required during zebrafish neurogenesis ^35,38^. The neuromuscular deficiencies observed in *dlx5a/6a* morphants may result from altered expression of *msx1b* and cell adhesion molecules in premigratory NCCs. In mouse, *Dlx5* and *Msx2* genes regulate anterior neural tube closure through expression of EphrinA5-EphA7 involved in cell adhesion ^33^. This suggests that a common genetic network implicating *Dlx, Msx* and cell adhesion molecules is involved in neural plate border and neural crest cells during mouse and zebrafish neurulation.

These new data give insights for a better understanding of the cellular and molecular processes that could be altered in some human congenital NTDs, such as craniorachischisis that originates from defects of both anterior and posterior neurulations. The anterior and posterior NTDs observed in *Dlx5/6* mutant mice are also associated with hypospadias, characterized by midline urethral malformations, and limb ectrodactyly ^17,31,32^. Expression of *Dlx5/6* in the cloacal membrane, linked to the ventral ectodermal ridge ^59^, and in the genital tubercle is necessary for urethral formation. Moreover, it has been shown that *Dlx5/6* expression in the apical ectodermal ridge is required for proper appendage formation in mouse and zebrafish ^17,18,32^. The limb phenotype of *Dlx5/6*^*-/-*^ mice resembles that of patients with congenital split hand-foot malformation type I (SHFM-I), linked to genomic deletion or rearrangement in the *DLX5/DLX6* cluster locus. Altogether, the data unveil the role of *Dlx5/6* in ectodermal cells for the proper development of the neural tube, the urogenital system and limbs. Interestingly, it has been reported that NTDs can be associated with limb malformations and other midline defects, including urogenital and diaphragmatic disorders, as observed in Czeizel-Losonci syndrome ^61-65^. Our data bring new light on common aetiology for a spectrum of idiopathic anomalies characterizing certain human congenital disorders.

## METHODS

### Ethical statement

All experiments with zebrafish were performed according to the guidelines of the Canadian Council on Animal Care and were approved by the University of Ottawa animal care committee (institutional licence #BL 235 to ME). All efforts were made to minimize suffering; manipulations on animals were performed with the anaesthetic drug tricaine mesylate (ethyl 3-aminobenzoate methanesulfonate; Sigma-Aldrich, Oakville, ON, Canada). Embryos were killed with an overdose of the latter drug.

Procedures involving mice were conducted in accordance with the directives of the European Community (council directive 86/609) and the French Agriculture Ministry (council directive 87–848, 19 October 1987, permissions 00782 to GL).

### Animal maintenance

Zebrafish and their embryos were maintained at 28.5°C according to methods described in ^66^. Wild-type adult zebrafish were kept and bred in circulating fish water at 28.5°C with a controlled 14-h light cycle. Wild-type, controls, and injected embryos were raised at similar densities in embryo medium in a 28.5°C incubator. Embryos were treated with 0.0015% 1-phenyl 2-thiourea (PTU) to inhibit melanogenesis. Embryos were killed with an overdose of tricaine mesylate for analysis.

Mice were housed in light, temperature (21°C) and humidity controlled conditions; food and water were available ad libitum. WT animals were from Charles River France. The mouse strain *Dlx5/6*^*+/-*^ ^25,27,28,32^ was maintained on a hybrid genetic background resulting from crosses between C57BL/6N females and DBA/2JRj males (B6D2N; Janvier Labs, France).

### Morpholino-mediated knockdown

The morpholino-mediated knockdown was validated as previously described ^18^. We performed injection or co-injection of *dlx5a* and/or *dlx6a* MOs in 1 cell-stage embryos (0.4 mM or 0.8 mM). The 5’-untranslated region of *dlx* genes has been used to design translation-blocking antisense MOs against *dlx* transcripts. The following translation-blocking MOs were obtained from Gene Tools (LLC, Philomath, OR, USA): *dlx5a* MO 5’-TCCTTCTGTCGAATACTCCAGTCAT-3’; *dlx6a* MO 5’-TGGTCATCATCAAATTTTCTGCTTT-3’.

Splice-blocking MOs targeting exon 2 excision were also designed to confirm the phenotype obtained using the translation-blocking MOs: *dlx5a* e2i2 MO 5’-TATTCCAGGAAATTGTGCGAACCTG-3’; *dlx6a* e2i2 MO 5’-AAATGAGTTCACATCTCACCTGCGT-3’ (from Gene Tools, LLC).

As controls, we injected water or 1.6 mM of Standard Control MO (Gene Tools) that targets a human beta-globin intron mutation that causes beta-thalessemia. (Gene Tools 5’-CCTCTTACCTCAGTTACAATTTATA-3’). To ensure MOs specificity, rescue of morphant phenotypes was performed by co-injecting the corresponding *dlx5a/dlx6a* transcripts mutated on the MO target site (*dlx5a* MO binding site T(ATG)ACTGGAGTATTCGACAGAAGGA, *mutdlx5a* sequence C(ATG)ACGGGTGTTTTTGATAGGAGGA; *dlx6a* MO binding site AAAGCAGAAAATTTG(ATG)ATGACCA, *mutdlx6a* sequence ATTGCAAATAATATG(ATG)ATGACCA). Mutated transcripts were synthesized using the SP6 mMessage mMachine kit (Ambion). The quantitative analysis of control, knockdown and rescue experiments are detailed in ^18^. Based on experimental results, co-injection of *dlx5a/dlx6a* MOs (0.8 mM each) and resulting severe phenotype specimens were selected for analysis.

### Histological analyses

*In situ* hybridization on whole-mount zebrafish embryos were performed as previously described ^18^.

For whole-mount immunostaining on zebrafish embryos, dechorionated embryos were fixed in 4% paraformaldehyde (PFA) in 1X phosphate buffered saline (PBS) overnight at 4°C, washed in PBST (PBS 0.1% Tween), dehydrated in methanol and stored in methanol 100% at - 20°C. The samples were then rehydrated in a graded methanol-PBST series and treated with PBDT (PBS 1% DMSO 0.1% Tween). Cells were immunodetected on wholemount embryos with mouse anti-SV2 monoclonal antibody diluted in PBDT an incubated overnight at 4°C (1:100, DHBS). After 5 rounds of 30 min washes in PBST, embryos were incubated overnight at 4°C with secondary anti-mouse HRP-conjugated antibody diluted in PBST (1:200, Jackson Immuno), washed with 5 times 30 min in PBST and revealed with DAB chromogenic substrate.

For immunostaining on cryosections, foetuses were fixed 3h in 4% PFA 0.5 % Triton X- 100 at 4°C, washed overnight at 4°C in PBST, cryopreserved in 30% sucrose in PBS and embedded in OCT for 12-16μm sectioning with a Leica cryostat. Cryosections were dried for 30 min and washed in PBS. Rehydrated sections were blocked for 1h in 10% normal goat serum, 3% BSA, 0.5% Triton X-100 in PBS. Primary antibodies were diluted in blocking solution and incubated overnight at 4°C (Tnnt3 antibody, 1/200, T6277, Sigma; Tuj1, 1/1000, BLE801202, Ozyme). After 3 rounds of 15 min washes in PBST, secondary antibodies were incubated in blocking solution 2h at RT together with 1μg/ml Hoechst 33342 to visualize nuclei. Secondary antibodies consisted of Alexa 488 or 555 goat anti-rabbit or anti-mouse isotype specific (1/500, Jackson Immunoresearch). After 3 rounds of 15 min washes in PBST, slides were mounted in 70% glycerol for analysis.

Skeletal preparations on mouse foetuses were performed as previously described ^67^.

## DATA AVAILABILITY

Availability of materials and data upon request

## Supporting information

## ACKNOWLEDGMENTS

We thank Joachim Wittbrodt for the donation of the zebrafish ncad and ncam3 plasmids. A particular thank goes to the team in charge of animal care, Vishal Saxena, Stéphane Sosinsky and Fabien Uridat, and to Pr. Amaury de Luze in charge of animal well-being. We thank Mss. Aicha Bennana and Lanto Courcelaud for administrative assistance. We also thank Dr. Benoit Robert for helpful discussions.

This research was partially supported by the EU Consortium HUMAN (EU-FP7-HEALTH-602757), the ANR grants TARGETBONE (ANR-17-CE14-0024) and METACOGNITION (ANR-17-CE37-0007) to NNN/GL and by CIHR grant MOP137082 to ME. EH was a recipient of a postdoctoral fellowship from the government of Canada.

## AUTHOR CONTRIBUTIONS

GL and EH conceived and designed the study. NNN and EH carried out the experiments. ME and GL provided materials. EH wrote the original manuscript. All authors discussed the results and contributed to manuscript revision.

