## Supplementary material for "Posterior axis formation requires *Dlx5/Dlx6* expression at the neural plate border"

**Supplementary Figures**

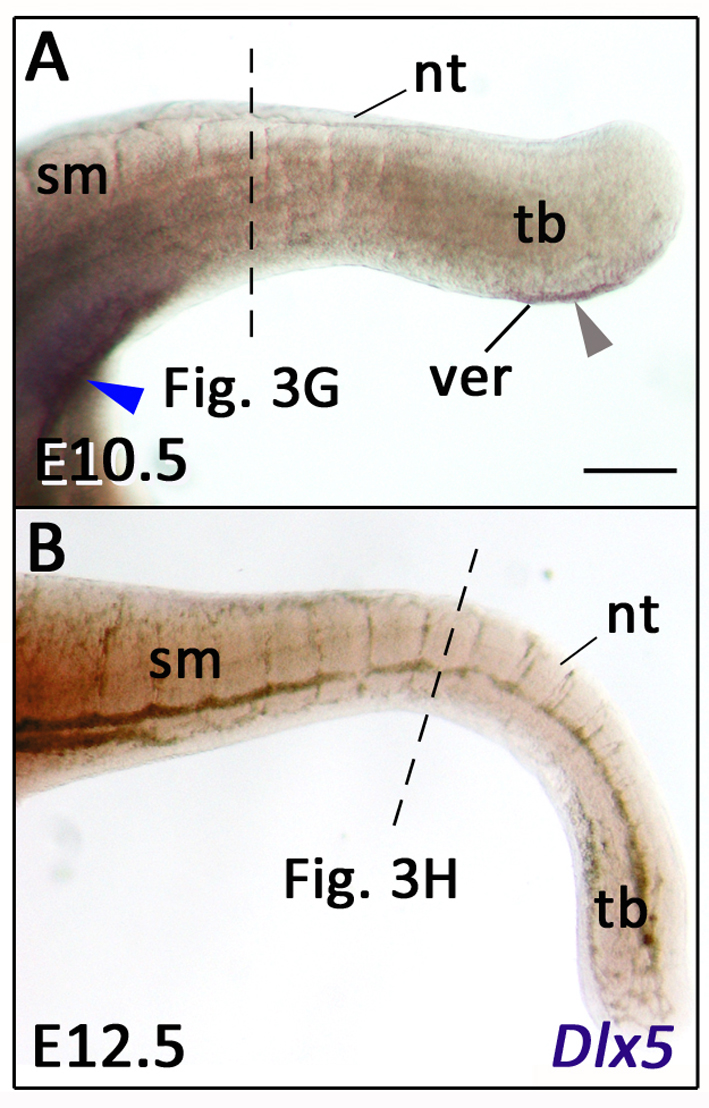

**Supplementary Fig. S1. *Dlx5* expression after posterior neuropore closure in mouse.**

(A, B) Lateral views of whole-mount *in situ* hybridization for *Dlx5* in mice at E10.5 and E12.5. The dashed lines indicate the section levels analysed in Fig. 3G-H. *Dlx5* expression is detected at the cloacal level and in the VER at E10.5 (A, blue and grey arrowheads respectively) but is not detectable at E12.5. Abbreviations: nt, neural tube; sm, somitic mesoderm; tb, tail bud; ver, ventral ectodermal ridge. Scale bar in A for A-B 200µm.

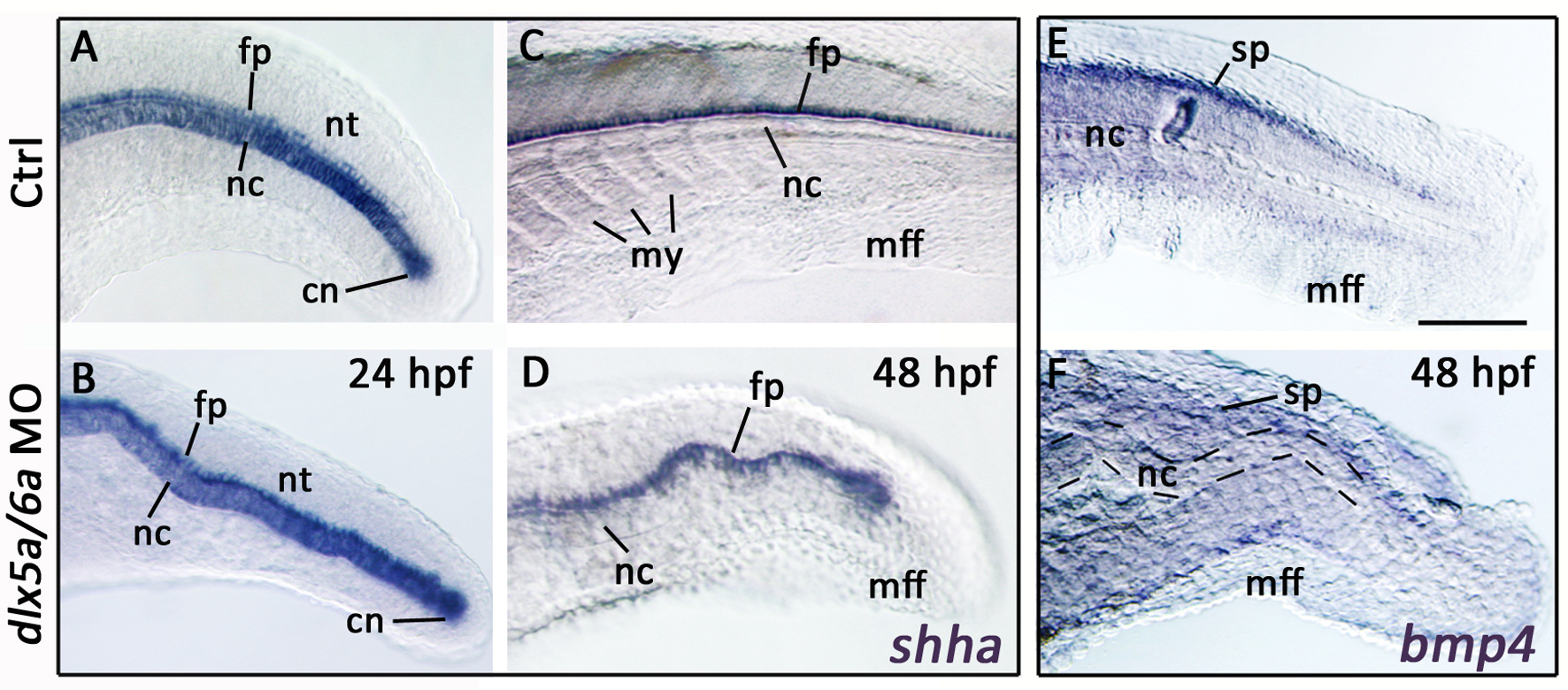

**Supplementary Fig. S2. Expression of *shha* and *bmp4* during zebrafish spinal neurulation.**

(A-F) Lateral views of whole-mount *in situ* hybridization in controls and *dlx5a/6a* zebrafish morphants for *shha* at 24 hpf and 48 hpf (A-D) and for *bmp4* at 48 hpf. The expression of *shha* in the notochord and the neural tube floor plate well reveal the ondulating axis phenotype observed in *dlx5a/6a* morphants (A-D). At 48 hpf, the axis malformation is associated with a decrease of *bmp4* expression in the spinal cord (E-F). Abbreviations: cn, chordoneural hinge; fp, floor plate; mff, median fin fold; my, myotomes; nc, notochord; nt, neural tube; sp, spinal cord. Scale bar in E for A-F 100µm.

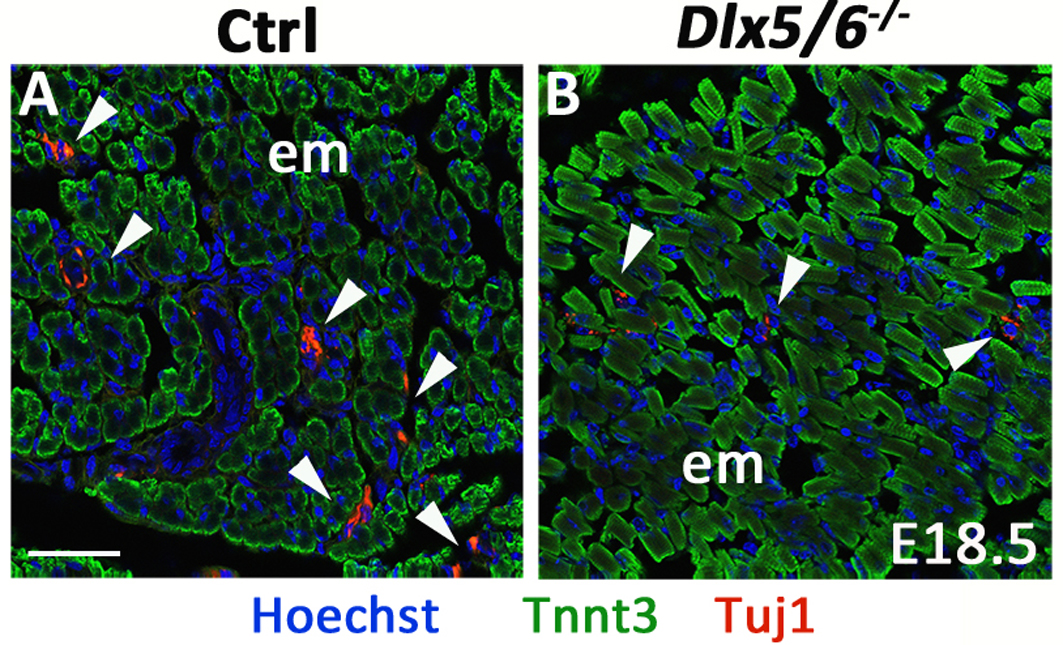

**Supplementary Fig. S3. Defect of neuromuscular innervation in *Dlx5/6^-/-^* mice.**

(A-B) Immunostaining on coronal cryosections for Tnnt3 and Tuj1 in epaxial muscles of control and *Dlx5/6^-/-^* foetuses at E18.5. The *Dlx5/6^-/-^* mutants show defect of neuromuscular innervation in epaxial musculature. Abbreviations: epm, epaxial muscles. Scale bar in A for A-B 20µm.
